# Proofreading deficiency in mitochondrial DNA polymerase does not affect total dNTP pools in mouse embryos

**DOI:** 10.1101/2020.03.04.971283

**Authors:** Sushma Sharma, Camilla Koolmeister, Phong Tran, Anna Karin Nilsson, Nils-Göran Larsson, Andrei Chabes

**Author notes:** ARISING FROM Hämäläinen et al. *Nature Metabolism* https://doi.org/10.1038/s42255-019-0120-1 (2019).

## Abstract

Mitochondrial DNA (mtDNA) mutator mice express proofreading-deficient mtDNA polymerase gamma (Polg^D257A^) and age prematurely^1,2^, whereas mice with other mitochondrial defects do not show global signs of early aging. The reason for this discrepancy is not completely understood. Hämäläinen *et al.* recently reported that both induced pluripotent stem cells (iPSCs) and primary mutator embryonic fibroblasts harboring Polg^D257A^ demonstrate increased mtDNA replication frequency leading to depletion of the total cellular dNTP pool^3^. The authors proposed that the decreased availability of cellular dNTPs for nuclear genome replication leads to compromised nuclear genome maintenance and premature aging. Here we report that total cellular dNTP pools are normal in mtDNA mutator mouse embryos (genotype: *Polg*^D257A/D257A^), which shows that a living organism exclusively expressing Polg^D257A^ has normal total dNTP pools despite ubiquitous rapid cell division.

## Introduction

In actively proliferating eukaryotic cells, dNTPs are produced primarily *de novo* in the cytosol and transported into the nucleus and mitochondria for DNA synthesis. The dNTP pool balance in dividing cells is tightly controlled by the allosterically regulated enzymes ribonucleotide reductase and dCMP deaminase^4^, and even minor dNTP pool imbalances are mutagenic^5,6^.

The allosteric regulation of eukaryotic ribonucleotide reductase, which has two separate allosteric sites, is one of the most sophisticated among all enzymes ever studied and is highly conserved among different species^7^. The allosteric *activity* site determines the overall dNTP concentration. When this site binds ATP, the enzyme is stimulated, and when it binds dATP the enzyme is inhibited. The allosteric *specificity* site determines the dNTP pool balance, and it binds four different allosteric effectors. Binding of dATP and ATP stimulates the production of dCTP and dTTP, binding of dTTP stimulates the production of dGTP, and binding of dGTP stimulates the production of dATP. dCMP deaminase balances the levels of dCTP and dTTP and is allosterically stimulated by dCTP and inhibited by dTTP. Because ribonucleotide reductase and dCMP deaminase are highly abundant during S phase in mitotic cells, both the overall dNTP pool and the balance of the individual dNTPs are continuously fine-tuned by these enzymes.

dNTPs can also be catabolized in the cytosol to corresponding deoxynucleosides (dNs) either by stepwise dephosphorylation, in which the last step of conversion of dNMPs to dNs is carried out by 5’-nucleotidases, or by one-step dephosphorylation by the dNTP triphosphohydrolase SAMHD1, another enzyme that is allosterically controlled by dNTPs^4^. dNs can be transported into the mitochondria where dedicated mitochondrial deoxyribonucleoside kinases phosphorylate them to dNMPs that in turn are phosphorylated to form dNTPs. dNs can also be transported between cells and similarly phosphorylated to dNTPs through salvage pathways^4^. Salvage pathways play a more important role in dNTP pool maintenance in non-dividing cells, in which *de novo* dNTP production is very low.

dNTP pools are determined either by enzymatic assays that utilize incorporation of radiolabeled nucleotides by DNA polymerases, as in Hämäläinen *et al.*, or by HPLC-based separation coupled with UV or mass-spectrometry detection. The enzymatic assays have several drawbacks. First, metabolites present in cell extracts might affect DNA polymerase activity. In particular, this concerns NTPs that are present in logarithmically growing cells in concentrations exceeding those of dNTPs by ~30- to ~170-fold depending on the NTP/dNTP pair^8^. NTPs are to a certain extent misincorporated by DNA polymerases and thus compete with dNTPs in the assay^9^. The use of certain DNA polymerases minimizes interference from NTPs^10^, but probably does not completely eliminate it. Second, the concentrations of the four dNTPs are measured in four separate assays, which adds variability to the results. Third, the enzymatic assays lack an internal standard that can be measured in the same reaction. We note that Hämäläinen *et al.* provide “relative dNTP levels” for iPSCs and mitochondria in Figure 2 and “[total dNTPs] µM” for the embryonic fibroblasts in Extended Data Figure 3a, but they do not explain how they standardized the reported dNTP levels or how they calculated the molar concentrations of dNTPs in embryonic fibroblasts, for which a careful measurement of cellular volume is required. We also note that the reported concentrations of dNTPs (between 0.1 and 0.3 µM) are 50- to 200-fold lower compared to what is established in the field^11^. Such low cellular dNTP concentrations would not support processive DNA replication.

HPLC-based assays using strong anion-exchange separation coupled with UV-detection avoid these drawbacks. First, NTPs do not interfere with the detection of dNTPs because they are removed by a boronate affinity step, and most other metabolites in the cell extracts are eluted early during the HPLC separation^12^. Second, all four dNTPs are measured during the same HPLC separation, eliminating possible variations due to sample handling. Third, an aliquot of the cell extract prior to NTP removal can be used as an internal standard for correction of possible losses during the extraction procedure and for normalization of the amount of material used for the preparation of extracts.

## Results

We measured dNTP and NTP levels in embryonic day (E)13.5 mouse *Polg*^D257A/D257A^ embryos because at this mid-gestation period there is massive growth and the embryo consists primarily of actively dividing mitotic cells in their natural milieu. We found no decrease in the total dNTP pools in the homozygous *Polg*^D257A/D257A^ embryos compared to wild-type and heterozygous embryos (Fig 1a). Furthermore, in contrast to the results of Hämäläinen *et al.*, there were no differential decreases in dATP and dTTP. Our representative HPLC chromatograms show nearly identical levels of dNTPs and NTPs in embryos of all three genotypes (Fig 1 b and c). As a positive control to show the sensitivity of our method for measuring nucleotide pool balances, we measured dNTP and NTP pools in *Samhd1*-knockout E13.5 embryos, which are known to have increased dNTP pools^13^. As expected, dNTP pools were elevated in *Samhd1*-knockout E13.5 embryos compared to wild-type embryos by 1.5-fold to 4.5-fold (Fig 1 a, c, and d). The greatest increase was for the dGTP pool, which is in agreement with previously published data^13,14^.

**Figure 1.**
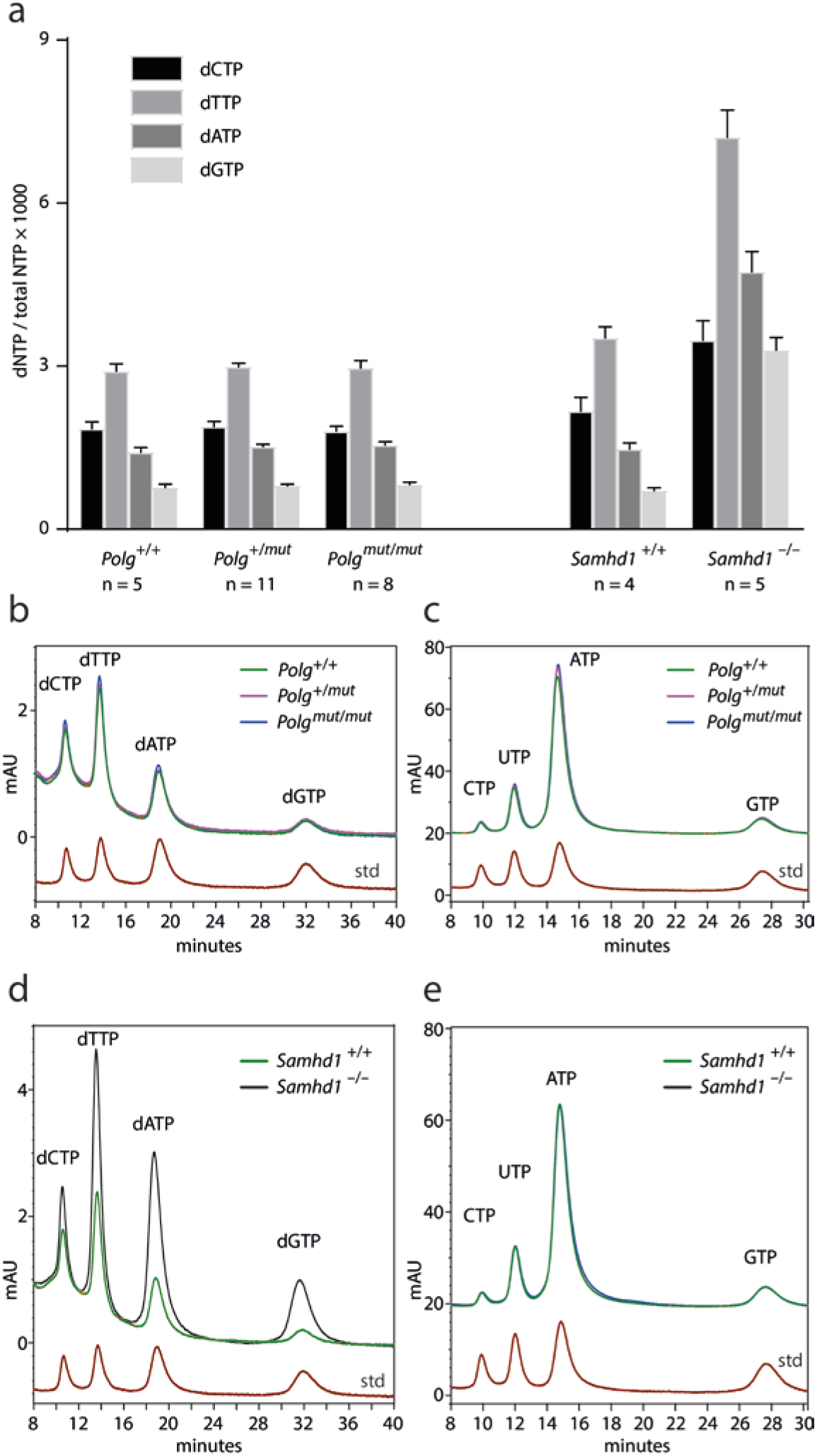
dNTP pools in E13.5 mouse embryos. **a**, Quantification of dNTP pools in wild-type (*Polg*^+/+^), heterozygous Polg^D257A^ (*Polg*^*+/mut*^), and homozygous Polg^D257A^ (*Polg*^*mut/mut*^) E13.5 mouse embryos shows no difference among the three genotypes. Quantification of dNTP pools in wild-type Samhd1 (*Samhd1*^*+/+*^) and Samhd1 homozygous knockout (*Samhd1*^*−/−*^) E13.5 mouse embryos as positive controls. n = number of embryos. Data are presented as mean ± SD. **b**, An overlay of representative HPLC chromatograms of the OD_260_ absorbance of dNTPs in the *Polg*^*+/+*^, *Polg*^*+/mut*^, and *Polg*^*+mut/mut*^ embryos. **c**, An overlay of the corresponding HPLC chromatograms of the OD_260_ absorbance of NTPs in the *Polg*^*+/+*^, *Polg*^*+/mut*^, and *Polg*^*+mut/mut*^ embryos used in **b**. **d**, An overlay of representative HPLC chromatograms of the OD_260_ absorbance of dNTPs in the wild-type and *Samhd1*^*−/−*^ embryos. **e**, An overlay of the corresponding HPLC chromatograms of the OD_260_ absorbance of NTPs in the wild-type and *Samhd1*^*−/−*^ embryos used in **d**. The dNTP and NTP standards are shown in red.

We conclude that mtDNA mutator embryos with rapid ubiquitous cell division have normal dNTP pools and thus that changes in dNTP pools cannot explain the premature aging phenotype.

## Methods

To isolate Polg^1^ and Samhd1^14^ embryos of different genotypes, female mice were paired individually overnight with males and inspected for vaginal plugs the next morning. At E13.5, the pregnant females were euthanized by cervical dislocation. The embryos were immediately dissected out in ice-cold PBS and the tails were taken for genotyping. The embryos were immediately snap-frozen in Eppendorf tubes in liquid nitrogen and stored at −80°C. After the addition of ice-cold 12% (wt/vol) trichloroacetic acid, 15 mM MgCl_2_ solution, and glass beads (1 mm Zirconia/Silica beads, BioSpec), the embryos were thawed on ice and homogenized on a BeadBeater (BioSpec) for 2 × 30 s at 4 °C in a cold room. The supernatant was collected by centrifugation at 20,000 × *g* for 5 min at 4 °C and neutralized with an ice-cold mixture of 98% trioctylamine and 10 ml Freon as described in^12^. Aliquots (30 µl) were saved for analyses of NTPs, and the rest of the aqueous phase was pH adjusted with 1 M ammonium carbonate (pH 8.9), loaded onto a boronate column (Affi-Gel Boronate Gel, Bio-Rad) and eluted with 50 mM ammonium carbonate (pH 8.9) and 15 mM MgCl_2_ to separate dNTPs from NTPs. The eluates containing dNTPs were adjusted to pH 3.4 and analyzed on a LaChrom Elite® HPLC system (Hitachi) with a Partisphere SAX HPLC column (Hichrome, UK). NTPs were analyzed on the same column using 30 µl aliquots of the aqueous phase adjusted to pH 3.4.

## Author contributions

C.K. isolated and genotyped Polg mouse embryos and P.T. and A.K.N. isolated and genotyped Samhd1 mouse embryos. S.S. performed the dNTP pool measurements. S.S., N.-G.L., and A.C. analyzed the data. A.C. wrote the manuscript.

